# The relative importance of host phylogeny and dietary convergence in shaping the bacterial communities hosted by several Sonoran Desert *Drosophila* species

**DOI:** 10.1101/2024.05.31.596909

**Authors:** James G. DuBose, Thomas Blake Crook, Luciano M. Matzkin, Tamara S. Haselkorn

**Affiliations:** Department of Biology, University of Central Arkansas, Conway, Arkansas, USA; Department of Biology, Emory University, Atlanta, Georgia, USA; Department of Entomology, University of Arizona, Tucson, Arizona, USA; BIO5 Institute, University of Arizona, Tucson, Arizona, USA; Department of Ecology and Evolutionary Biology, University of Arizona, Tucson, Arizona, USA

## Abstract

Complex eukaryotes vary greatly in the mode and extent that their evolutionary histories have been shaped by the microbial communities that they host. A general understanding of the evolutionary consequences of host-microbe symbioses requires that we understand the relative importance of host phylogenetic divergence and other ecological processes in shaping variation in host-associated microbial communities. To contribute to this understanding, we described the bacterial communities hosted by several *Drosophila* species native to the Sonoran Desert of North America. Our sampling consisted of four species that span multiple dietary shifts to cactophily, as well as the dietary generalist *D. melanogaster*, allowing us to partition the influences of host phylogeny and extant ecology. We found that bacterial communities were compositionally indistinguishable when considering incidence only but varied when considering the relative abundances of bacterial taxa. Variation in community composition was not explained by host phylogenetic divergence but could be partially explained by dietary variation. In support for an important role of diet as a source of ecological selection, we found that specialist cactophilic *Drosophila* deviated more from neutral predictions than dietary generalists. Overall, our findings provide insight into the evolutionary and ecological factors that shape host-associated microbial communities in a natural context.

## Introduction

Numerous studies have documented substantial variation in the microbial communities associated with hosts from across the tree of life. In some cases, the variation between host-associated microbial communities mirrors the phylogenetic divergence between the hosts themselves (Brooks et al. 2016; Groussin et al. 2017). Paired with several examples of functionally important symbioses, this has led to the idea that microbial influences on host evolution are ubiquitous (McFall-Ngai et al. 2013; Bordenstein and Theis 2015). However, it is becoming increasingly apparent that not all complex multicellular organisms host resident microbial communities (Hammer et al. 2019), highlighting that hosts vary in the extents that their evolutionary histories have been shaped by microbial symbioses. Despite vast interest in the evolutionary consequences of host-microbe symbioses, we still lack a general understanding of what aspects of host natural history and extant ecology facilitate more or less persistent interactions with microbial symbionts.

The term *phylosymbiosis* was recently proposed to describe patterns of divergence between host-associated microbial communities that parallel the patterns of host phylogenetic divergence (Brooks et al. 2016; Lim and Bordenstein 2020; Trevelline et al. 2020). The most commonly suggested explanation for phylosymbiotic patterns is host-selection for or against certain microbial populations within their associated communities (Bordenstein and Theis 2015). For example, different species within the genus *Hydra* express distinct antimicrobial peptides to shape their associated bacterial communities, which is thought to have greatly influenced their diversification (Franzenburg et al. 2013). More recent appreciation has been given to the simpler explanation that phylosymbiotic patterns can emerge from more passive filtering caused by host physiological (e.g. gut pH) and ecological variation (e.g. diet) that tracks their phylogeny (Mazel et al. 2018). However, it is frequently suggested that variation in host ability to selectively assemble their microbial communities has been an important driver of evolutionary change in complex eukaryotes (Bordenstein and Theis 2015; Brooks et al. 2016). While this might be the case in some instances, patterns of phylosymbiosis can also be explained by other aspects of host ecology and microbial community assembly dynamics that are not related to host fitness or host evolutionary divergence (Moran and Sloan 2015; Mazel et al. 2018; Kohl 2020). Therefore, finding support for or against a phylosymbiotic signal does not inherently lend insight into the underlying processes involved in generating said pattern.

An important conceptual advance in community ecology describes the processes that drive variation in ecological communities as analogous to those that drive variation in allele frequencies: selection, drift, dispersal (migration), and speciation (diversification more generally) (Vellend 2010). Each of these processes can act in tandem with varying degrees of effect. Understanding the relative importance of these process in shaping community variation is necessary to understand the contexts in which we might expect microbial communities to have significantly influenced host evolution. For example, the phylosymbiosis model suggests that host associated microbial communities are deterministically assembled in concordance with host phylogeny. However, the relative importance of deterministic and stochastic (drift) processes varies. The consistency that a host-associated microbial community can be deterministically assembled and remain stable can be weakened by increased sensitivity to drift (Vega and Gore 2017; Mallott and Amato 2021; Taylor and Vega 2021). Although this has clear implications for when to expect patterns of phylosymbiosis and how to interpret them, studies that have documented co-divergence between hosts and their microbial communities make little or no effort to examine the relative importance of stochastic and deterministic processes in generating said pattern (Brooks et al. 2016; Trevelline et al. 2020). Likewise, dispersal and diversification can also play significant roles in shaping variation in communities. For example, in the absence of mechanisms to homogenize dispersal, hosts that sample different pools of microbes are expected to have different microbial communities that reflect (at least in part) the diversification of microbes within their respective environments.

Previous studies of the relationship between host phylogeny and the assembly of their microbial communities have focused on host selection while attempting to control for other sources of variation (Brooks et al. 2016; Trevelline et al. 2020). However, host factors that are not correlated with their phylogenetic divergence can potentially have a greater influence on their microbial community assembly relative to host selection (Muegge et al. 2011; Sullam et al. 2012; Mallott and Amato 2021). Therefore, understanding the potential for microbial communities to have influenced their host’s evolutionary history also requires an understanding of the relative importance of alternative selective drivers. Furthermore, laboratory studies on the relevance of microbial communities and key taxa to their hosts often show inconsistent results with and across taxa, illustrating a high degree of context-dependency (Douglas 2018; Mallott and Amato 2021). While these studies are necessary for making more causal inferences, it is important to ground their findings in the host population’s natural context. To contribute to a more general understanding of what aspects of host natural history and extant ecology shape their interactions with microbial symbionts, here we described the bacterial communities hosted by several *Drosophila* species located in the Sonoran Desert of North America. Our sampling consisted of four species that span multiple dietary shifts to cactophily, as well as the dietary generalist *D*. *melanogaster*. This sampling allowed us to quantify relative importance of a possible source of selection that is not tightly correlated with host phylogeny (Markow and O’Grady 2008). We included the bacterial phylogeny in our quantification of community composition to assess the potential for bacterial diversification in the environment to drive community variation between hosts. After quantifying the relative importance of host phylogeny and diet in shaping community variation, we used a neutral model to infer and contrast the relative importance of stochastic and deterministic processes in shaping observed variation between hosts of different dietary guilds.

## Materials and Methods

### Drosophila collection

Our objective was to compare the relative importance of host phylogeny and diet in shaping their bacterial communities. Therefore, we sampled five sympatric *Drosophila* species with dietary variation that is not tightly correlated with their phylogenetic divergence. Cactophilic specialists, which feed on one cactus species, included *D. mettleri*, *D. nigrospiracula*, and *D. mojavensis.* We also included *D. arizonae*, a cactus generalist that feeds on multiple cactus species and occasionally other food, as well as *D. melanogaster*, which is a dietary generalist and serves as a phylogenetic outgroup. *D. mojavensis* and *D. arizonae* are closely related sister species, diverging 840,000 years ago (Benowitz et al. 2022). *D. mojavensis* feeds on the Organ Pipe cactus *Stenocereus thurberi* in the Sonoran Desert and *D. arizonae* is a cactus generalist (Fellows and Heed 1972; Heed 1978; Ruiz and Heed 1988; Matzkin 2014). In contrast, *D. nigrospiracula* and *D. mettleri* are more distantly related, but both feed on the Saguaro cactus *Carnegiea gigantea* in the Sonoran Desert. Likewise, *D. melanogaster* and *D. arizonae* are distantly related but both exhibit more generalist diets. Natural population samples of *D. arizonae, D. melanogaster, D. mettleri, D. mojavensis,* and *D. nigrospiracula* were collected from banana-baited traps in October 2019. *D. arizonae* (n=10) and *D. melanogaster* (n=5) were collected from Tucson, Arizona, while *D. mettleri* (n=6), *D. mojavensis* (n=5), and *D. nigrospiracula* (n=7) were collected from Organ Pipe National Monument, Arizona.

### DNA extraction and sequencing

DNA extraction on individual flies were carried out using the DNeasy blood and tissue kit (Qiagen) using the standard protocol with the following modifications. To ensure that the bacteria extracted are from inside the fly (primarily gut bacteria), each sample was sterilized by a full vortex in 200β*L* of 50% bleach for 2 minutes, followed by three full vortexes in 200β*L* of sterile water at 1 minute each. The samples were then homogenized with a sterilized pestle in a small test tube containing 200β*L* of enzymatic lysis buffer, and then incubated at 37°C for 90 minutes. 200β*L* of AL buffer and 20β*L* of proteinase K were added to the samples, and another incubation was performed at 56°C for 90 minutes, and the rest of the protocol was done according to the Qiagen DNeasy kit protocol. Samples were stored at -80°C before being sent to Novogene (Sacramento, CA) for 16S rRNA sequencing using the 314F and 806R primers.

### Sequence Processing

Sequence demultiplexing, as well as removal of adapter, primer, and low-quality sequences was initially performed by Novogene using their standard quality control pipeline. All subsequent sequence processing steps were implemented using QIIME2 v.2023.2 (Bolyen et al. 2019). We used the Divisive Amplicon Denoising Algorithm (DADA2) pipeline to trim low quality bases from the ends of reads, perform additional chimeric sequence removal, and merge paired reads (Callahan et al. 2016). To assign taxonomy to each sequence, we first used the REference Sequence annotation and CuRatIon Pipeline (RESCRIPt) (Robeson et al. 2021) to curate the SILVA SSU NR99 database (v. 138.1), thus improving sequence classification. Briefly, the database was dereplicated, the V3-V4 regions of each sequence were extracted, and the database was dereplicated again. The q2-feature-classifier plugin (Bokulich et al. 2018) was then used to train an amplicon-specific Naïve Bayes classifier on the curated database, as well as assign taxonomic classifications to each amplicon sequence variant (ASV). ASVs that were classified as either mitochondria or chloroplast, as well as ASVs that did not have at least an 80% alignment identity across 80% of any reference sequence in the SILVA database, were discarded. Finally, we constructed a phylogenetic tree by inserting the quality-controlled sequences into the SILVA reference database using SATe-enhanced phylogenetic placement (Mirarab et al. 2011), and all fragments that were not able to be inserted were discarded from further analyses.

### Bacterial community diversity and dissimilarity analyses

After inspecting rarefaction curves for each sample, all samples were rarefied to a depth of 19,000 sequences, which was approximately the maximum depth that allowed all samples to be retained. We evaluated differences in microbial community diversity (alpha-diversity) by comparing ASV richness, Shannon index, Pielou’s evenness, and Faith’s phylogenetic diversity across sample groups, and statistical differences were determined using Kruskal-Wallis tests with Benjamini-Hochberg false discovery rate adjustments and an alpha of 0.05. We evaluated differences in microbial community composition (beta-diversity) by comparing weighted and unweighted UniFrac distances between sample groups (Lozupone and Knight 2005; Lozupone et al. 2007). Permutational multivariate analyses of variance (PERMANOVA) were used to evaluate differences between groups and permutational multivariate analyses of dispersion (PERMDISP) were used to test for equality of variances between groups, each with 999 permutations, Benjamini-Hochberg false discovery rate adjustment, and an alpha of 0.05. All aforementioned diversity metrics and statistics were calculated using the q2-diversity plugin (Bolyen et al. 2019).

### Quantifying evidence of phylosymbiosis

To quantify the relationship between host phylogenetic divergence and dissimilarity in their bacterial communities, we first constructed a host phylogeny based on partial sequences (∼660 base pairs) of the cytochrome oxidase subunit 1 gene that were generated through amplification using the LCO 1490 and HCO 2198 universal CO1 primers (Supplemental Table 1). Sequences were retrieved from the NCBI nucleotide database and aligned using the MUSCLE algorithm (Edgar 2004). We then used IQ-TREE (Nguyen et al. 2015) to generate a maximum likelihood phylogenetic tree with a GTR+F+G4 substitution mode (selected based on lowest Bayesian Information Criteria). We then constructed hierarchical clustering graphs based on weighted and unweighted UniFrac distances, and visually compared tree topologies.

To calculate the correlation between host phylogenetic divergence and bacterial community dissimilarity, we first calculated the pairwise phylogenetic distances between each species using the *cophenetic* function from the *ape* R package (Paradis and Schliep 2019). We then conducted a Mantel test using the host phylogenetic distance matrix and UniFrac distance matrices (weighted and unweighted). Mantel tests were implemented using the *mantel* function from the *vegan* R package (Oksanen et al. 2022).

### Partitioning variance in bacterial communities between host phylogeny and diet

To quantitatively assess the contributions of host relatedness and diet in driving the compositional variation across bacterial communities, we used distance-based redundancy analyses (dbRDA). To vectorize phylogenetic distances so that they could be used in the dbRDA model, we defined phylogenetic distance as each species’ distance from *D. melanogaster* (which also serves as an outgroup). To calculate how much variance in weighted UniFrac and unweighted UniFrac distances could be explained by phylogenetic distance and dietary differences, which were categorically defined as organ pipe cactus, saguaro cactus, and generalist, we performed a distance-based redundancy analysis using sthe *dbrda* function from the *vegan* R package (Oksanen et al. 2022). Statistical significance of each regression model considered in the distance-based redundancy analysis was assessed by using the *anova* base R function to perform permutational multivariate analyses of variance with 999 permutations (R Core Team 2022). We then partitioned the amount of variance explained by diet and phylogenetic distance using the *varpart* function from the *vegan* R package (Oksanen et al. 2022).

### Modeling neutrally assembled bacterial communities

To identify how consistent the bacterial distributions across *Drosophila* gut communities were with neutral dynamics, we modelled the frequency of each bacterium (ASV) under stochastic birth, death, and immigration processes using the neutral community model proposed by Sloan et al., 2006. Briefly, we defined the regional pool of bacteria as the average relative abundance of each bacterium across all samples, and assume that all bacteria are equally capable of dispersing to each local community (*Drosophila* gut). Thus, the probability a randomly lost individual is replaced from the regional pool is proportional to its abundance within the regional pool times an estimated immigration rate that is assumed to be equal for all bacterial taxa. Assuming all bacterial taxa have equal fitness within a given local community (*Drosophila* gut), the relationship between the frequency at which a bacterial taxon occurs can be modeled as a function of its abundance in the regional pool using a beta distribution. Therefore, if the frequency of a given ASV can be predicted by its abundance in the regional pool, its distribution is consistent with neutral dynamics.

To test if the observed community was consistent with neutral expectations, we fit the observed ASV frequencies and relative abundances to those predicted under neutrality using nonlinear least-squares regression. In addition to constructing a neutral model for the total ASVs across all bacterial communities, we constructed separate models for generalists and specialists, where the regional pool was redefined accordingly. All model construction and fitting was done using the *tyRa* R package (Sprockett 2018).

## Results

### Diversification between Drosophila-associated bacterial communities is not evident

Our goal was to quantify the variation between *Drosophila*-associated bacterial communities and gain insight into the processes involved in generating said variation. Diversification between the members of different communities has the potential to drive patterns of variation in community composition. Given the relatively short life span of *Drosophila*, potential bacterial diversification would likely occur in the environmental pools of bacteria that colonize their guts. To include an estimate of bacterial diversification in quantifying compositional dissimilarity, we calculated unweighted UniFrac distances between each bacterial community (Lozupone and Knight 2005). We then used a PERMANOVA to test for compositional differences and a PERMDISP to test for differences in the extent of compositional turnover across individuals in a given species. This showed that the *Drosophila*-associated bacterial communities were not phylogenetically distinct (PERMANOVA: F=1.17, p=0.132; PERMDISP: F=1.03, p=0.393) (Figure 1A).

**Figure 1.**
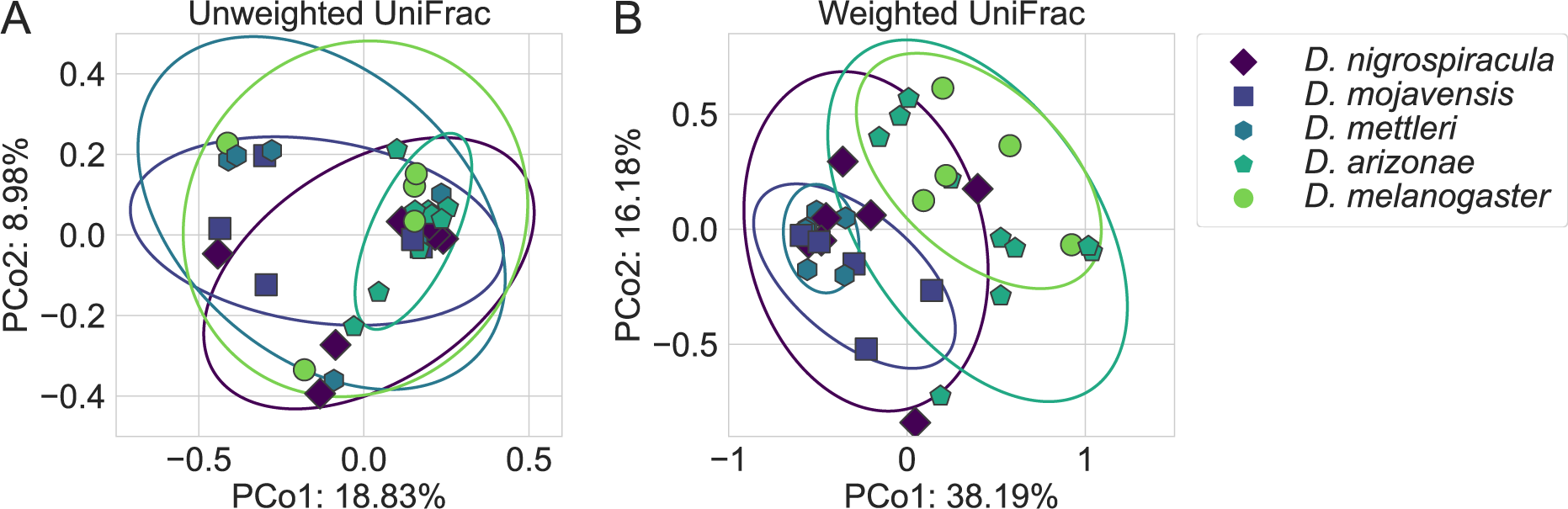
The bacterial communities associated with the sampled *Drosophila* hosts do not vary in composition (A) but vary in structure between generalists (*D. melanogaster* and *D. arizonae*) and cactus specialists (*D. mojavensis, D. mettleri,* and *D. nigrospiracula*) (B). In both principal component analysis plots, each point represents a bacterial community, and points that are closer to each other are more similar. A) Principal component analysis was performed based on unweighted UniFrac distances, which considers community composition and phylogeny, but not relative abundances. B) Principal component analysis was performed using weighted UniFrac distances, which considers community composition and phylogeny, as well as the relative abundance of each bacterial population. In each plot, ellipses represent two standard deviation confidence intervals.

### Drosophila-associated bacterial communities vary in composition when considering the relative abundance of each taxa

To include the relative abundances of bacterial taxa in our quantification of community composition, we calculated the weighted UniFrac distance between each bacterial community. In addition to incorporating the bacterial phylogeny, weighted UniFrac includes the relative abundance of each ASV in calculating community dissimilarity (Lozupone et al. 2007). This showed that bacterial communities varied in composition across host species (F=3.37, p=0.001), which could not be attributed to differences in dispersion (F=1.97, p=0.085) (Figure 1B). Pairwise comparisons showed that the two more generalist flies (*D. melanogaster* and *D. arizonae*) did not differ from each other (F=0.54, p=0.729) but differed from all other specialist cactophilic flies (F=3.323 – 11.638, p ≤ 0.0183). Likewise, the bacterial communities hosted by specialist cactophilic flies were not compositionally differently, with the exception of *D. mettleri* and *D. nigrospiracula* (F=2.05, p=0.0271). Full quantifications can be found in Supplemental Table 2.

We then examined the potential for alpha-diversity to have contributed to the patterns of variation in community composition. For example, two communities with the same set of bacteria could show compositional variation if one community is more even than the other. Overall, we did not find differences in alpha-diversity when considering ASV richness (H=5.27, p = 0.262), Shannon diversity (H=3.39, p = 0.495), Pielou’s evenness (H=0.112, p = 0.250), or Faith’s phylogenetic diversity (H=5.39, p = 0.116) (Figure 2).

**Figure 2.**
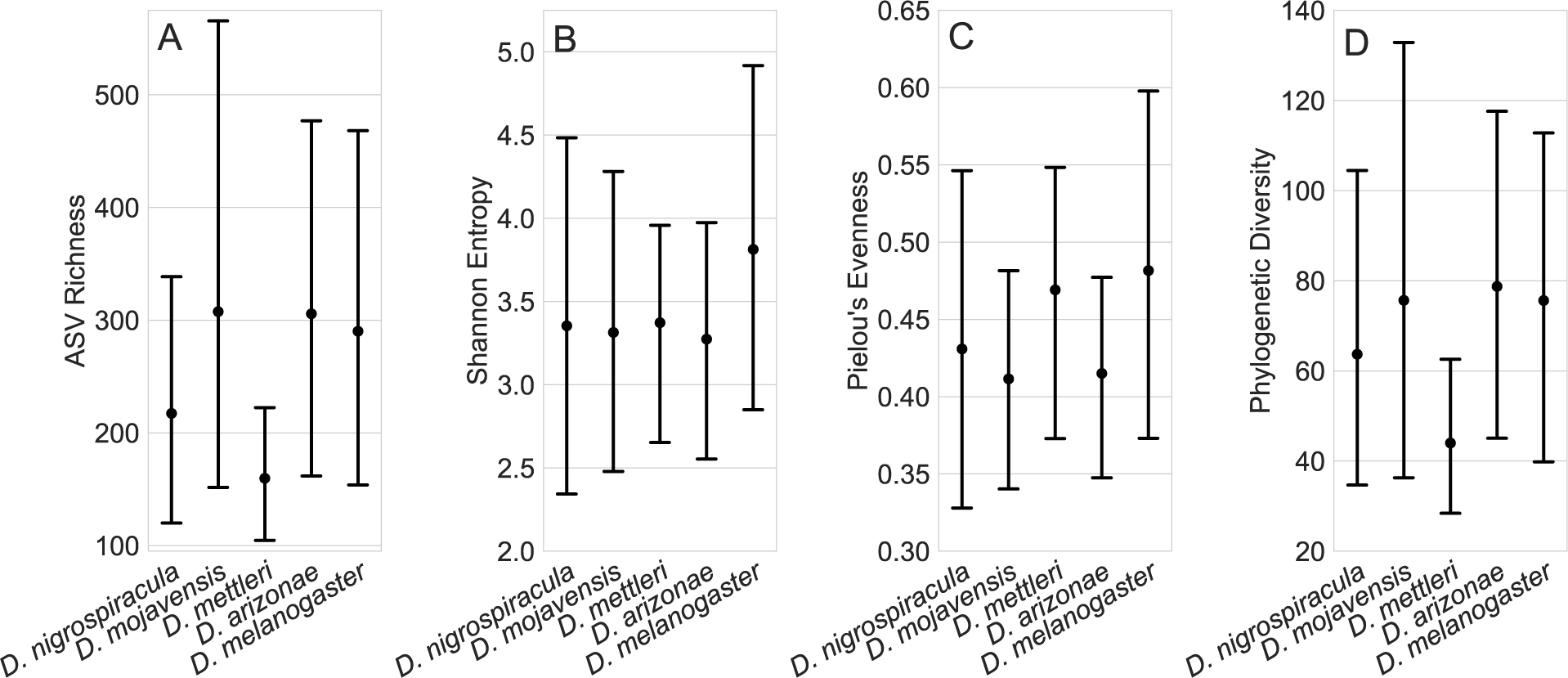
The bacterial communities associated with the sampled *Drosophila* hosts do not vary in alpha-diversity, regardless of host identity or dietary guild. Point plots show the *Drosophila* gut bacterial community alpha diversity as measured using A) ASV richness, B) Shannon entropy, C) Pielou’s evenness, and D) Faith’s phylogenetic diversity. Points indicate mean and error bars indicate 95% confidence intervals.

### Patterns of dissimilarity between bacterial communities do not correspond to their host’s phylogenetic divergence

Deterministic assembly in concordance with host phylogeny is one possible contributor to the aforementioned variation in bacterial community composition. Although observing a pattern consistent with phylosymbiosis doesn’t lend insight into the processes involved in generating said pattern, quantifying a phylosymbiotic signal is a useful first step in a more detailed analysis. We first constructed hierarchical clustering graphs based on unweighted and weighted UniFrac dissimilarities and examined their congruence with the host phylogeny (Figure 3A and B). This showed that divergence in bacterial community composition did not correspond with patterns of host phylogenetic divergence when considering incidence only (Figure 3A) or relative abundances (Figure 3B). These results are consistent with our previous analysis, where there were not differences in community composition when considering incidence only, and variation in community composition when considering relative abundances appeared related to diet and not host relatedness. To quantitively test for a pattern of phylosymbiosis, we used a Mantel test to see how well host phylogenetic divergence predicted dissimilarity in their bacterial communities. This showed no correlation between host phylogeny and bacterial community composition when considering incidence only (r_M_ = 0.042, p=0.183) (Figure 3C) or relative abundances (r_M_ = 0.067, p=0.122) (Figure 3D).

**Figure 3.**
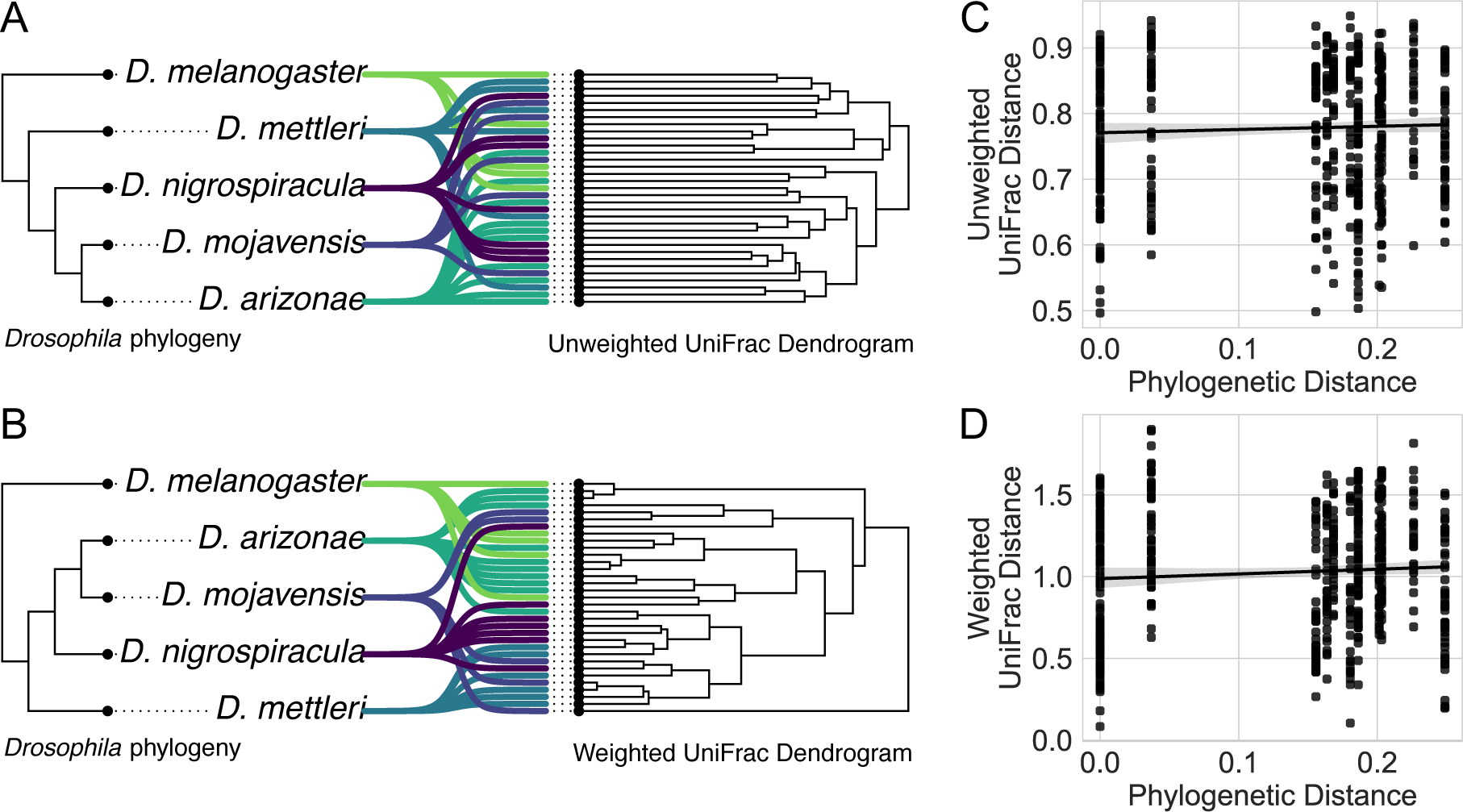
Bacterial community composition and structure do not vary with host phylogeny. Tree congruence plots show the relationship between the sampled *Drosophila* phylogeny and dissimilarity of the bacterial communities, as measured by A) unweighted UniFrac and B) weighted UniFrac. Tree congruence plots show the *Drosophila* phylogeny on the left and dendrogram of their associated bacterial communities on the right, and each line represents an individual bacterial community. Overall, there is a lack of topological congruence between the *Drosophila* phylogeny and their bacterial community composition and structure. C and D show the correlation between phylogenetic distance between hosts and dissimilarity between their community composition (unweighted UniFrac) and structure (weighted UniFrac) respectively. If more distantly related hosts had less similar bacterial communities, these plots would show negative correlations. However, no correlation is apparent.

### Variation in bacterial community composition can be partially explained by host diet but not relatedness

The previous analysis suggested that the sampled bacterial communities do not vary in concordance with host phylogeny. However, we did find that the bacterial communities hosted by more generalist *Drosophila* were dissimilar from those hosted by cactophilic specialist *Drosophila* (Figure 1B). To quantify how much variation in community composition could be explained by host diet and relatedness, we used a distance-based redundancy analysis and variance partitioning. This showed that host diet explained 22.14% of the variance in community composition when considering relative abundances (F = 5.38, p < 0.001) (Figure 4). However, host relatedness did not explain a significantly explain any variation (F = 1.08, p = 0.344). Consistent with our finding that bacterial community composition did not significantly differ between *Drosophila* species when considering incidence only, we found that neither host diet (F = 0.132, p = 0.076) nor phylogeny (F = 1.034, p = 0.394) explained a significant portion of the variation in community composition (Figure 4).

**Figure 4.**
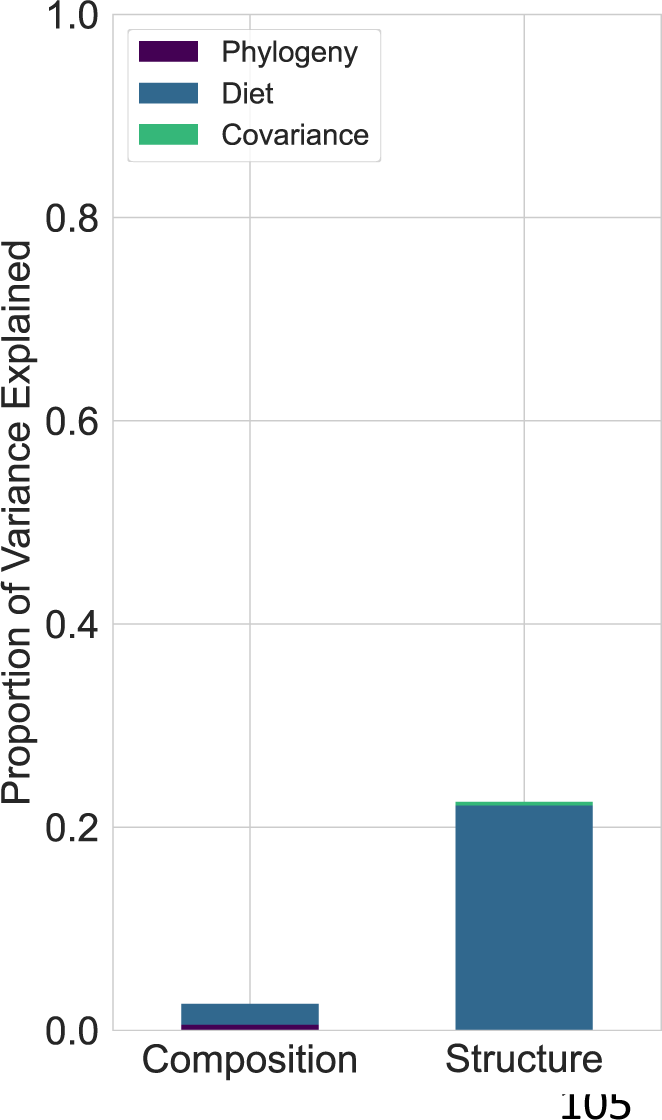
A portion of variation in bacterial community structure between hosts can be explained by host dietary differences, but not host phylogeny. The stacked bar plot shows the proportion of variation in bacterial community composition (unweighted UniFrac distances) and structure (weighted UniFrac distances) explained by host diet and phylogeny. Significant variation in community composition between hosts was not found, so the lack of explained variance was expected.

### The bacterial communities associated with cactophilic specialist Drosophila deviate more from neutral predictions than more generalist Drosophila species

After finding that dietary differences explain a reasonably large portion of the variation in bacterial community composition observed across hosts, we wanted to compare the relative importance of selective and neutral processes between dietary guilds. We used a model that predicts the prevalence of bacterial taxa across local communities (*Drosophila* hosts) as a function of their total relative abundance, assuming stochastic birth, death, and immigration. We then quantified how well the observed distribution of bacterial taxa matched the neutral expectations. Across all hosts, neutral predictions explained 49.46% of the variance in the observed distribution of bacteria across all *Drosophila*-associated communities (R^2^ = 0.4946) (Figure 5A). When examining only more generalist hosts, neutral predictions explained 52.3% of the variance in the distribution of bacteria across generalists (R^2^ = 0.523) (Figure 5B). In contrast, cactophilic hosts deviated significantly more from neutral predictions, which only accounted for 24.81% of the observed variance (R^2^ = 0.2481) (Figure 5C).

**Figure 5.**
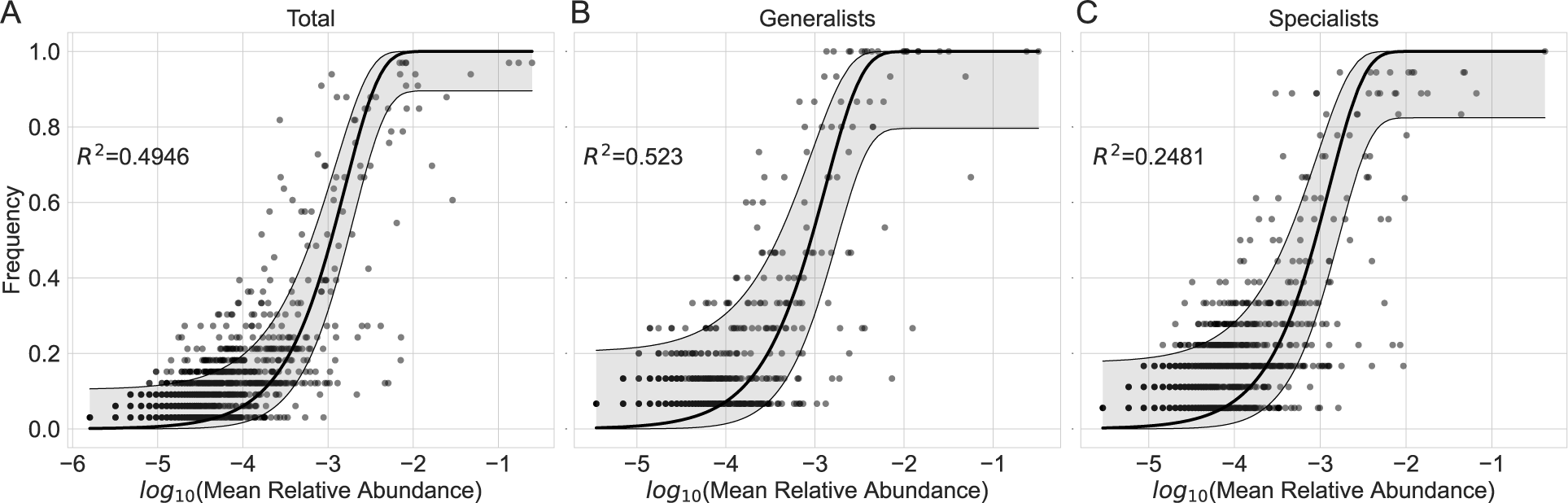
Plots showing the predicted frequency of bacteria across local communities (Drosophila guts) as a function of their relative abundance within a regional pool. Separate models were constructed for A) the total bacterial community across all sampled *Drosophila*, B) generalists only, and C) specialists only. For each plot, individual points indicate observed ASVs. The y-axis denotes the frequency (proportion of *Drosophila*) that an ASV was observed, and the x-axis denotes the log_2_ of the ASV’s average relative abundance across samples. The black line indicates the predicted values based on a neutral model, and the shaded gray area around said line indicates the 95% confidence interval.

## Discussion

Complex eukaryotes vary greatly in the mode and extent that their evolutionary histories have been shaped by the microbial communities that they host. A general understanding of the evolutionary consequences of host-microbe symbioses requires that we understand the relative importance of host phylogenetic divergence and other ecological processes in driving variation in host-associated microbial communities. In the five *Drosophila* species that we sampled in this study, we did not find evidence of extensive diversification between the sampled bacterial communities, leading to compositional similarity between *Drosophila* hosts when only incidence was considered (Figure 1A). However, we did find that bacterial communities varied in composition when considering relative abundances (Figure 1B), which could not be explained by differences in alpha diversity (Figure 2). We found that this variation in community structure was not related to host phylogenetic divergence (Figure 3), but that it could partially be explained by variation in *Drosophila* diet (Figure 4). Specifically, cactophilic specialists and more generalist *Drosophila* species hosted bacterial communities that were occupied by similar sets of bacteria that varied in their relative abundances across host species. In support for an important role of diet in structuring the sampled *Drosophila*-associated communities, we found that cactophilic *Drosophila* deviated more from neutral predictions than more dietary generalists (Figure 5).

One of the most frequently proposed explanations for observed variation in host-associated microbial communities is that hosts select for or against specific bacterial taxa that are presumably relevant to their fitness. Over evolutionary time, such interactions are thought to generate patterns of phylosymbiosis: variation in host-associated microbial communities parallel host phylogenetic divergence (Bordenstein and Theis 2015; Lim and Bordenstein 2020). Our results were inconsistent with this pattern. A possible explanation for this lack of phylosymbiotic signal is phylogenetic scale. For example, the relationship between host phylogeny and similarities in their bacterial communities deteriorates in more distantly diverged mammalian lineages (Groussin et al. 2017). However, the timescale at which this pattern was examined across mammalian hosts was comparable to or greater than the divergence time between the *Drosophila* hosts sampled in this study. Therefore, we could expect that if host phylogeny was important, the bacterial communities hosted by the sister species *D. mojavensis* and *D. arizonae* would be compositionally similar, which we did not find support for. Furthermore, a lack of correlation between *Drosophila* relatedness and bacterial community dissimilarity has also been reported in other studies as well (Chandler et al. 2011; Adair et al. 2020). Taken together, these studies highlight inconsistencies between the functional roles of *Drosophila*-associated bacteria found in laboratory studies and their ecological relevancies indicated in natural populations. For example, laboratory manipulations have found that bacterial community composition can alter *D. melanogaster* life history strategies and drive adaptation (Rudman et al. 2019; Walters et al. 2020). While these findings have prompted some to suggest that *Drosophila*-associated bacterial communities have significant consequences for their host’s evolution, to our knowledge, signatures of such evolutionary dynamics have yet to be supported in surveys of natural *Drosophila* populations. Furthermore, it has been suggested that bacterial community differences do not influence reproductive isolation in *D. melanogaster* (Leftwich et al. 2017), and different studies have found contradictory effects of specific bacteria on *D. melanogaster* lifespan and physiology (Douglas 2018; Fast et al. 2018). These discrepancies illustrate the importance of informing laboratory-derived insight with natural studies.

More generally, several previous studies of how variation in host-associated microbial communities relates to host phylogeny attempted to control for alternative sources of ecological selection, such as diet and the environment (Brooks et al. 2016; Trevelline et al. 2020). However, accounting for, rather than controlling for, the effects of these alternative selective drivers is necessary to understand the evolutionary dynamics between hosts and their microbial communities. Here we focused on diet, and in contrast to the lack of correlation with host phylogeny, we found dietary variation was able to explain a significant portion of variation in community structure between host species. This suggests that differences in diet could select for or against different bacterial population. However, it is important to note that dietary differences alone do not explain most of the variation in community structure seen across hosts. Given the exploratory nature of this study, our inferences about the processes involved in generating the structural variation between more generalist- and specialist cactophilic-associated communities are limited.

Bacterial communities associated with specialist cactophilic *Drosophila* deviated more from neutral predictions than those associated with more generalists, suggesting a decreased importance of stochasticity that is consistent with an important role of diet as a selective mechanism. However, it is difficult to infer the relative importance of deterministic and stochastic processes without controlled experiments and time-series sampling, so we interpret these results lightly. Furthermore, neutral models assume equivalent fitness within and dispersal to local communities (*Drosophila* guts). Given the descriptive nature of this study, we cannot discern the relative importance of these processes. We can however note that we did not find evidence of extensive diversification between the communities of different hosts. Given the relatively short life span of *Drosophila*, potential bacterial diversification would likely occur in the environmental pools of bacteria that colonize their guts. Therefore, the observed lack of diversification between hosts suggests that the *Drosophila* species are sampling common pools of environmental bacteria that are capable of dispersing to each host. Some evidence that dispersal bias is important in shaping *Drosophila*-associated bacterial communities has been found in laboratory studies, where several bacterial populations hosted by *D. melanogaster* are constantly replenished through food consumption (Blum et al. 2013). However, natural surveys have found that the most abundant bacterial taxa in *Drosophila*-associated communities are often in low abundance or absent from their food, suggesting that the importance of dispersal variation in shaping *Drosophila* gut bacterial communities may vary by species and/or differ in natural contexts (Martinson et al. 2017*a*, 2017*b*). Alternatively, it is possible that the persistence of some bacterial populations is trait mediated, as the guts of many insects have structural and chemical features that some bacteria have evolved to better exploit (Engel and Moran 2013). Future studies would be needed to discern the importance of these processes in the *Drosophila* species sampled in this study.

Underlying the similarity in bacterial community composition across host species, we found that the majority of *Drosophila*-associated bacterial communities were comprised of relatively few bacterial taxa. These findings are consistent with many previous studies, and add to a growing body of evidence that many bacteria are filtered from holometabolous insect guts (Chandler et al. 2011; Wong et al. 2011; Hammer et al. 2017; Martinson et al. 2017*a*; Adair et al. 2018; Henry and Ayroles 2022). From the perspective of habitat suitability, the guts of holometabolous insects represent harsh environments for many potential bacterial colonists. Gut chemistry and restructuring throughout development introduces frequent disturbances that can reduce or eliminate bacteria on the gut surface, making it difficult to establish persistent populations (Moll et al. 2001). These frequent disturbances could contribute to the lack of a phylosymbiotic signal observed in this study and several others. If a given microbial community is not stable enough to provide consistent benefits (or detriments) to its host at a population level, it would not be expected to selectively influence the host population’s evolution. Overall, further work is needed to understand when and how microbial communities significantly influence their host’s evolution.

## Acknowledgements

We would like to thank Christopher P. Catano for helpful discussion on the framework and analyses implemented in this paper. We would like to thank the Arkansas Department of Higher Education Student Undergraduate Fellowship program and the University of Central Arkansas Southwest Energy Research Fellowship program for funding.

## Data availability

All 16S rRNA sequences generated to compare bacterial communities have been submitted to the Sequence Read Archive (SRA) under the accession numbers SRR25517041 through SRR25517073 and be accessed using the BioProject number PRJNA1000992. All code written for data analysis, as well as processed data (such as the ASV count table, neutral model predictions, sample metadata, etc.) can be accessed at hps://github.com/gabe-dubose/ncdmic.

## Appendix for

### 1 Methodological Summaries

#### 1.1 Cytochrome c oxidase subunit I accession numbers

**Table S1:**
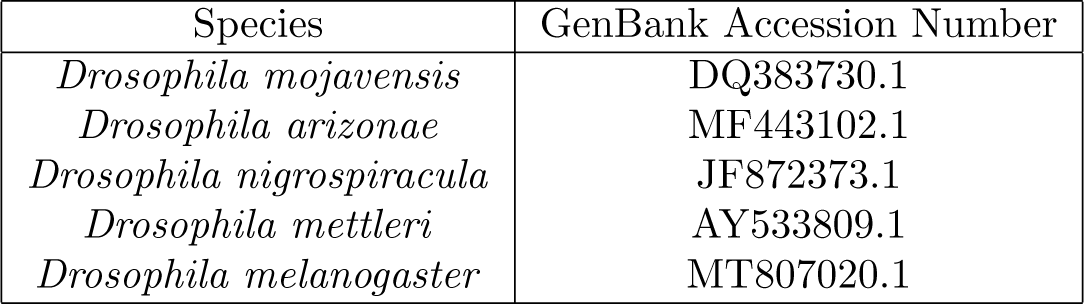
A table showing the the *Drosophila* species and GenBank accession number for the cytochrome C oxidase subunit I sequences used to infer the phylogeny of the sampled hosts.

| Species | GenBank Accession Number |
| --- | --- |
| *Drosophila mojavensis* | DQ383730.1 |
| *Drosophila arizonae* | MF443102.1 |
| *Drosophila nigrospiracula* | JF872373.1 |
| *Drosophila mettleri* | AY533809.1 |
| *Drosophila melanogaster* | MT807020.1 |

### 2 Supporting Results

#### 2.1 Taxonomic distributions across *Drosophila* hosts

**Figure S1:**
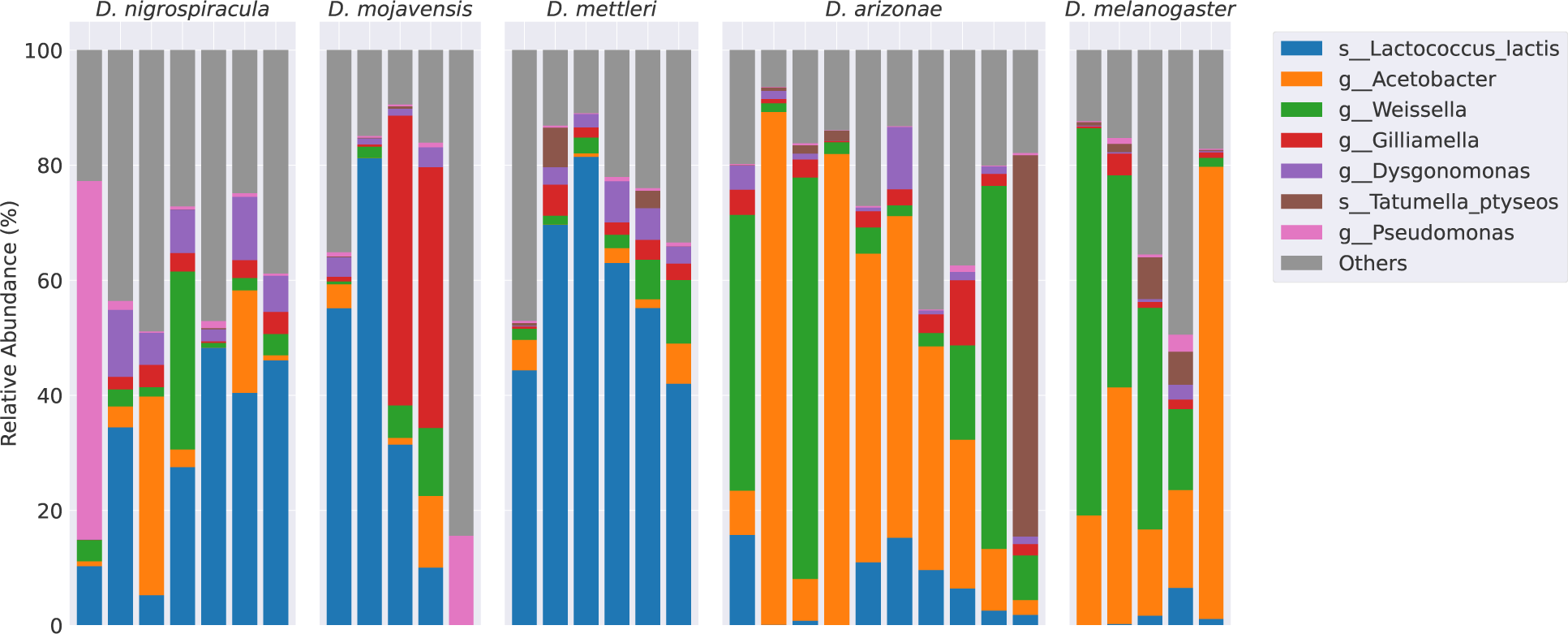
Taxonomy bar plots showing the relative abundances of the seven most abundant taxa found across the *Drosophila* species sampled in this study.

#### 2.2 Weighted UniFrac PERMANOVA results

**Table S2:** A table showing the full PERMANOVA results for compositional dissimilarity comparisons that were done quantified using Weighted UniFrac.

| Species 1 | Species 2 | F | p |
| --- | --- | --- | --- |
| *D.arizonae* | D.melanigaster | 0.540224 | 0.729 |
| *D.arizonae* | D.mettleri | 7.538250 | 0.015 |
| *D.arizonae* | D.mojavensis | 3.234842 | 0.0183 |
| *D.arizonae* | D.nigrospiracula | 3.685981 | 0.018 |
| *D.melanogaster* | D.mettleri | 11.637764 | 0.015 |
| *D.melanogaster* | D.mojavensis | 3.577240 | 0.018 |
| *D.melanogaster* | D.nigrospiracula | 3.945255 | 0.0167 |
| *D.mettleri* | D.mojavensis | 1.358943 | 0.276250 |
| *D.mettleri* | D.nigrospiracula | 2.046416 | 0.027143 |
| *D.mojavensis* | D.nigrospiracula | 0.645536 | 0.729 |

